# Synthesis and in vitro Activity of Eugenyl Benzoate Derivatives as BCL-2 Inhibitor in Colorectal Cancer with QSAR and Molecular Docking Approach

**DOI:** 10.1101/2020.08.17.253591

**Authors:** Fadilah Fadilah, Retnosari Andrajati, Ade Arsianti, Rafika Indah Paramita, Linda Erlina, Arry Yanuar

## Abstract

Eugenol derivatives can inhibit BCL-2 in HT29 colorectal cancer cells. This study is aimed to acquiring new compounds of Eugenyl benzoate (2-methoxy-4-(prop-2-en-1-yl)phenyl benzoate) derivatives that can inhibit HT29 colorectal cancer cells. In this research, we used several chemical reactions to synthesize novel compounds, such as Esterification, Demethylation, Halohydrin, and Sharpless reaction. Cytotoxicity assays were performed to test the inhibitory activity of compounds against HT29 colon cancer cells. QSAR analysis were carried out to analyse the relationship of chemical structure of the novel compounds with their cytotoxic activity. Ten novel compounds were successfully synthesized and tested *in vitro* against the HT29 cell. The IC_50_ of the novel compounds were between 26.56 μmol/ml - 286.81 μmol/ml which compound 4-[(2S)-2,3-dihydroxypropyl]-2-methoxyphenyl 2-hydroxybenzoate **(9)** showed as best active compound as BCL-2 inhibitors better than other synthesized compounds and Eugenol as well. QSAR analysis of the *in vitro* results gave a Log equation: 1/IC_50_ = −0.865-0.210 (LogP)^2^ + 1,264 (logP)-0.994 CMR (n = 10; r = 0.706; SE: 0.21; F = 0.497, sig = 7.86). The equation shows the log variable P and CMR affect IC_50_. The properties of hydrophobicity (log P) are more instrumental than the ones of steric (CMR).

## Introduction

According to the National Comprehensive Cancer Network (NCCN) guidelines, if the first-line and second-line treatments of colorectal cancer are unsuccessful, the therapy is targeted. Targeted therapy with the use of regorafenib is proven to be able to suppress the risk of death by 23%.[1] Monoclonal antibody-based therapies with EGFR target such as cetuximab and panitumumab in the metastatic phase (mCRC) are only effective 10-20% of mCRC due to molecular resistance mechanisms.[2] Cancer therapy, in addition to the above treatment, has also been performed through therapies using natural materials such as taxol, gossypol and other polyphenols.

Salicylic acid is used as a colorectal anticancer that can induce apoptosis and reduce the growth of SW480 colon cancer, HT-29 and HCT-116.[3] Another lead compound of salicylic acid used as a chemo preventive in colon cancer is shown to be safe to use in clinical studies and to reduce the growth of colon cancer in HT-29 cell.[4] The research of gallic acid can lead to the inhibition of HCT 15 colon cancer cells and induction of apoptosis in HT29 colon cancer cells.[5] Compounds of salicylic acid, salicylic acid and gallic acid have anticancer properties so that all three are condensed with eugenol in the position of the OH group. Another modification of eugenol derivatives in the allyl group through the halogen addition can increase the reactivity and specificity of the compounds. Compounds with electron-rich halogen groups are lipophilic and able to penetrate the lipid bilayer of the lead compounds. Besides that, the compounds can participate in molecular interactions that contribute to ligand-binding proteins.[6,7] An addition modification in the double bond group of Eugenol is performed by Hydroxylation and Halohydrin. The addition modification of the double bond of benzoate in the terminal hydroxyl group can increase anticancer activity as the existence of terminal hydroxyl group increases the coefficient of octanol-water partition, topological polar surface area (TPSA).[8] The HT29 cells of in vitro test are able to induce a mutated p53/Bax.[9] These cells in the study of Koehler *et al*. (2013) proved that over-expression of BCL-2 and BCL-xL occurred in the migration and invasion of colorectal cancer cells.

## Materials and Methods

### Materials

The tools used in the synthesis includes: ductless fume hood, refrigerators, technical and analytical balances, glassware, magnetic stirrers, chromatography and plates of KLT (Merck). The instruments used for structure elucidation include spectrophotometer IR (JASCO FT/IR-420 spectrophotometer), ^1^H-NMR spectroscopy (JEOL JNM-ECP500), ^13^C-NMR (JEOL JNM-ECP500), MS (Shimadzu GCMS QP-5000). The tools used in cytotoxicity test are Laminar Air Flow (LAF), micro pipettes of 200 −1000 μl, Eppendorf tube, and cell line well, talicytometer.

The materials used in the synthesis are Eugenol (Aldrich), ethanol p.a (Merck), absolute ethanol (MERCK), methanol p.a (Merck), acetone p.a (Merck), chloroform p.a (Merck), benzoyl chloride p.a (Merck), K2CO3 p.a (Merck), Na2CO3 p.a (Merck), Silica (Merck), sephadex L20 (Merck), TLC silica gel plate (MERCK), ethanol p.a (Merck), acetone p.a Merck, N-hexane p.a (Merck), ethyl acetate p.a (Merck), KBr Pro spectrophotometry, solvent for NMR (CDCl3). The materials used in the study include: Gallic acid (Sigma), Tetrahydrofuran (THF) (Wako), Diisopropylcarbodiimide (Sigma), DMAP (4-N,N-dimethylaminopyridine) (Wako), potassium carbonate (Wako), methyl iodide (Wako), Ethyl acetate, sodium bicarbonate (Wako), NaCl (Wako), MgSO4 anhydrous (Wako), chloroform (Merck), methanol, dimethylformamide (Wako), KHSO4 (Wako), LiOH monohydrate (Wako), hexanol (Wako).

The materials used in the in vitro assay are HT-29 cell line (The pathological anatomy lab of FMUI), aquadest, RPMI 1640 medium containing HEPES and GlutaMAX powder (Invitrogen Gibco BRL), MTT (Sigma), DMSO (Sigma), Fetal Bovine Serum/FBS (GIBCO), NaHCO3 (Sigma), solution of penicillin-streptomycin (GIBCO/ BTGIB), Phosphate Buffer Saline/PBS, 96-well plate, pipette tips (0, 2-1000 mL), filter unit 0.22 μm Millex GV (Millipore), trypan blue stain 0.25% (Gibco), Dextran Sodium sulfate/DSS (Sigma Chemical Company), Azoxymethane/AOM (Sigma Chemical Company), CMC Na (PT. Brataco). Materials for staining tissue include: Phosphate buffered formalin (E. Merck), high ethanol concentration (E. Merck), Xylol, paraffin block, Citrate Buffer, solution of hydrogen peroxide, antibody BCL-2 (Abcam, USA), PBS, N Universal (Wako), HRP-conjugated streptavidin (E. Merck), 3.3 ‘diaminobenzidine (DAB), Harris Hematoxylin.

### Methods

#### Brief synthetic procedure for the synthesis of targeted molecules

Eugenyl benzoate derivatives were synthesized using several chemical reactions, as seen in **Figure 1**. Compounds **(1-4)** were synthesized using esterification reaction between eugenol and salicylic acid, amino salicylic acid, gallic acid, mono methoxy gallic acid, respectively. Compound **(5)** and **(6)** were demethylated product of compound **(1)** and **(2)**, respectively. Furthermore, compound **(5)** and **(1)** were reacted with halohydrin reaction using N-chlorosuccinamide (NCS) and I2 as catalyst to give compound **(7)** and **(8)**, respectively. Compound **(1)** also used as starting material and reacted using (DHQ)2PHAL and (DHQD)2PHAL with Sharpless reaction that gave compound **(9)** and **(10)**, respectively.

**Figure 1.**
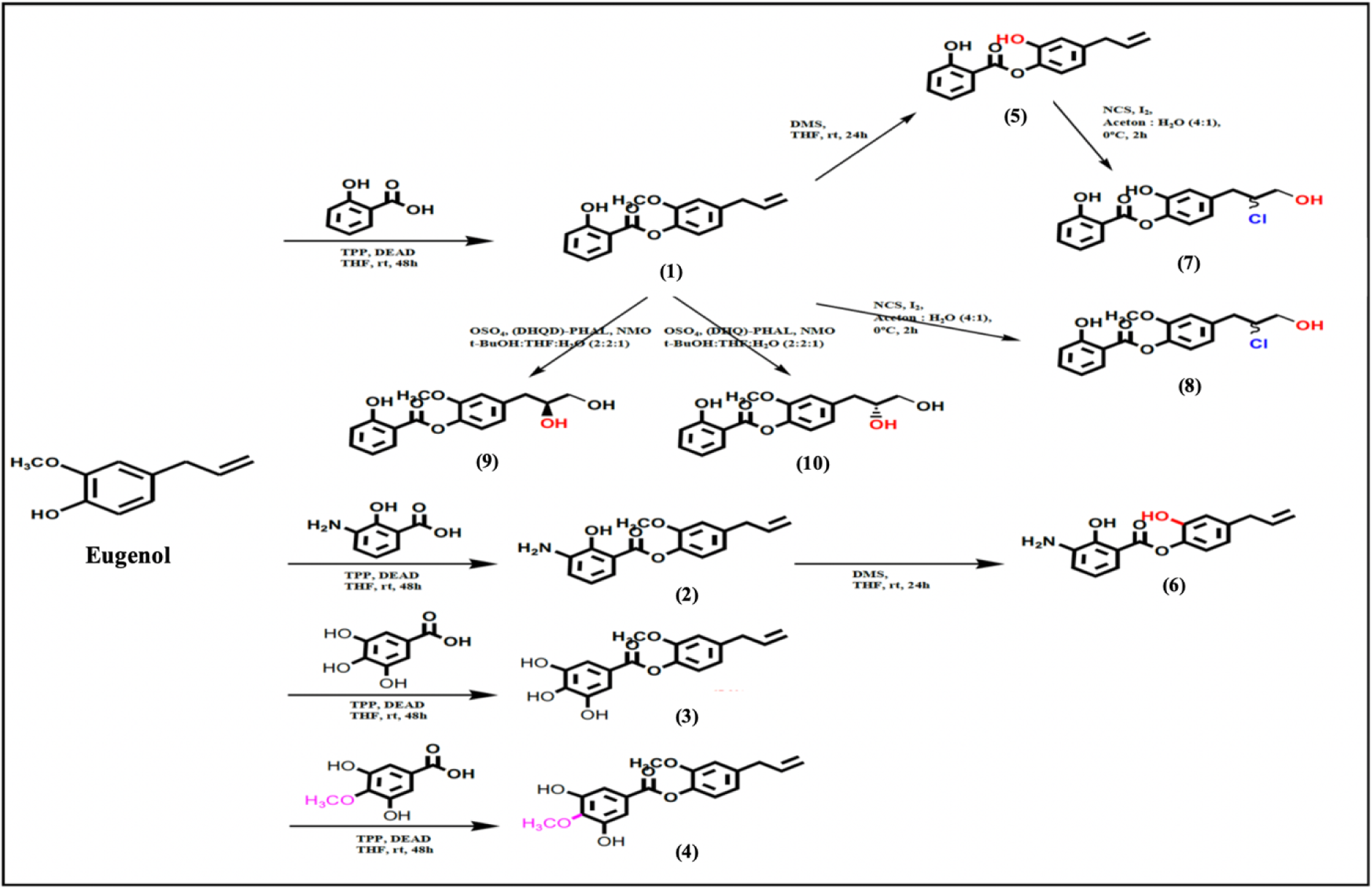
Synthesis scheme of Eugenyl benzoate derivatives

#### a. General procedure for preparation of compounds (1-4)

A solution of salicylic acid, gallic acid, or amino salicylic acid (1 mmol), respectively, eugenol (2 mmol) and 1.5 mmol diisopropylcarbadiimide (DIC) as catalysator in tetrahydrofuran (10 mL) were stirred in 0 °C for 30 min. In the mixture was then added solution of 0,1 mmol N,N-Dimethylaminopiridine (DMAP) in 1 ml THF and stirred at room temperature for 24 hours. The reaction mixture was then added aquadest and extracted with chloroform. The organic phase was added with MgSO_4_ anhydrate to take out the water residue. The product was purified using chromatography column with appropriate mobile phase. The spectral characters of synthesized compounds **(1-4)** are as follows.

### 2-methoxy-4-(prop-2-en-1-yl)phenyl 2-hydroxybenzoate (1)

White needle crystals, Yield: 75%, FTIR: 3403 (–OH), 2956 (C-H, aliphatic), 1683 (C=O), 1587 (C=C aryl), 1324 (C–O); ^1^H-NMR (500 MHz, DMSO-d6): δ 7.06 – 7.08 (d, 2H, =CH–, J= 8.0 Hz), δ 7.51 – 7.56 (m, 1H, =CH–), δ 8.11 – 8.12 (dd, 1H, =CH–, J= 8.0; 4.0 Hz), δ 6.84 – 6.85 (d, 2H, =CH–, J= 8.0 Hz), δ 7.02 – 7.04 (d, 1H, =CH–, J= 8.0 Hz), δ 3.40 – 3.42 (d, 2H, –CH2–, J= 6.9 Hz), δ 5.91 – 6.05 (m, 1H, =CH–), δ 5.08 – 5.19 (m, 2H, =CH2), δ 3.82 (s, 3H, –OCH_3_); ^13^C NMR (125 MHz, DMSO) d 168.5, 161.7, 151.5, 139.5, 137.0, 136.7, 136.0, 130.5, 122.4, 119.3, 116.2, 113.6, 111.8, 65.0, 40.0; MS m/z: 284.1 [M]^+^

### 2-methoxy-4-(prop-2-en-1-yl)phenyl 3-amino-2-hydroxybenzoate (2)

Light brown powder, Yield: 47.2%, FTIR: 3612 (–NH_2_), 3391 (–OH), 2460 (C-H, aliphatic), 1689 (C=O), 1440 (C=C aryl), 1137 (C–O); δ 6.83 – 6.84 (d, 2H, =CH–, J= 8.0 Hz), δ 7.58 – 7.59 (d, 1H, =CH–, J=7.8 Hz), δ 7.05 – 7.09 (d, 2H, =CH–, J= 8.0), δ 7.26 (s, 1H, =CH–), δ 3.40 – 3.42 (d, 2H, –CH_2_–, J= 6.9 Hz), δ 5.95 – 5.98 (m, 1H, =CH–), δ 5.10 – 5.12 (m, 2H, =CH_2_), δ 3.80 (s, 3H, –OCH_3_); ^13^C NMR (125 MHz, DMSO) d 169.1, 165.4, 156.7, 151.2, 137.7, 136.0, 135.4, 132.5, 122.4, 115.9, 113.5, 102.1, 100.9, 58.0, 40.0; MS m/z: 299.1 [M]^+^

### 2-methoxy-4-(prop-2-en-1-yl)phenyl 3,4,5-trihydroxybenzoate (3)

Blackish brown powder, Yield: 67.2%, FTIR: 3413 (–OH), 2950 (C-H, aliphatic), 1656 (C=O), 1527 (C=C aryl), 1388 (C–O); δ 7.*02* (s, 2H, =CH–), δ 6.80 – 6.82 (m, 2H, =CH–), δ 7.39 – 7.41 (d, 1H, =CH–, J=6.0 Hz), δ 3.39 – 3.41 (d, 2H, –CH_2_–, J= 6.0 Hz), δ 5.98 – 6.00 (m, 1H, =CH–), δ 5.10 – 5.12 (m, 2H, =CH_2_), δ 3.83 (s, 3H, –OCH_3_); ^13^C NMR (125 MHz, DMSO) d 165.4, 151.2, 145.2, 140.2, 137.7, 136.0, 135.4, 125.3, 122.4, 115.9, 113.5, 109.6, 58.0, 40.0; MS m/z: 316.0 [M]^+^

### 2-methoxy-4-(prop-2-en-1-yl)phenyl 3,5-dihydroxy-4-methoxybenzoate (4)

Yellowish powder, Yield: 59.1%, FTIR: 3372 (–OH), 2971 (C-H, aliphatic), 1698 (C=O), 1454 (C=C aryl), 1213 (C–O); δ 7.*02* (s, 2H, =CH–), δ 3.78 (s, 1H, –OCH_3_), δ 6.78 – 6.80 (m, 2H, =CH–), δ 7.37 – 7.39 (d, 1H, =CH–, J=6.0 Hz), δ 3.38 – 3.40 (d, 2H, –CH_2_–, J= 6.0 Hz), δ 5.96 – 5.98 (m, 1H, =CH–), δ 5.08 – 5.10 (m, 2H, =CH_2_), δ 3.81 (s, 3H, –OCH_3_); ^13^C NMR (125 MHz, DMSO) d 165.4, 152.0, 151.2, 145.2, 137.7, 136.0, 135.4, 125.3, 122.4, 115.9, 113.5, 109.6, 60.8, 58.0, 40.0; MS m/z: 330.1 [M]^+^

#### b. General procedure for preparation of compounds (5-6)

Dimethyl sulfide (2.5 mL) was added dropwise to a suspension of anhydrous AlCl_3_ (0.330 g, 2.5 mmol) in CH_2_Cl_2_ (5 mL) at 0 °C while stirring until AlCl_3_ was completely dissolved. Then the solution of compound **(1)** and **(2)**, respectively, (1.0 mmol) in CH_2_Cl_2_ (3 mL) was added in a period of 10 min at the same temperature. The mixture was allowed to stand at room temperature and stirred for 24 h. All dissolution process and the reaction were carried out in a nitrogen gas atmosphere. Afterward, 15 mL cold HCl 1 N was added and the mixture was extracted with CH_2_Cl_2_ (2×15 mL). The organic layer was dried with anhydrous MgSO_4_, filtered, and concentrated. The product was purified using chromatography column with appropriate mobile phase.[11] The spectral characters of synthesized compounds **(5-6)** are as follows.

### 2-hydroxy-4-(prop-2-en-1-yl)phenyl 2-hydroxybenzoate (5)

Light brown powder, Yield: 73.4%, FTIR: 3421 (–OH), 2884 (C-H, aliphatic), 1683 (C=O), 1415 (C=C aryl); ^1^H-NMR (500 MHz, DMSO-d6): δ 7.26 (d, 2H, =CH–, J= 8.0 Hz), δ 7.32 – 7.33 (m, 1H, =CH–), δ 8.08 – 8.11 (dd, 1H, =CH–, J= 8.0; 4.0 Hz), δ 6.61 – 6.63 (d, 2H, =CH–, J= 8.0 Hz), δ 6.78 – 6.80 (d, 1H, =CH–, J= 8.0 Hz), δ 3.26 – 3.28 (d, 2H, –CH2–, J= 6.9 Hz), δ 5.89 – 5.95 (m, 1H, =CH–), δ 5.10 – 5.11 (m, 2H, =CH2); ^13^C NMR (125 MHz, DMSO) d 168.5, 161.7, 151.5, 139.5, 137.0, 136.7, 136.0, 130.5, 122.4, 119.3, 116.2, 113.6, 111.8, 40.0; MS m/z: 270.3 [M]^+^

### 2-hydroxy-4-(prop-2-en-1-yl)phenyl 3-amino-2-hydroxybenzoate (6)

Dark brown semisolid, Yield: 58.3%, FTIR: 3620 (–NH_2_), 3430 (–OH), 2437 (C-H, aliphatic), 1656 (C=O), 1440 (C=C aryl); δ 6.83 – 6.84 (d, 2H, =CH–, J= 8.0 Hz), δ 7.58 – 7.59 (d, 1H, =CH–, J=7.8 Hz), δ 7.05 – 7.09 (d, 2H, =CH–, J= 8.0), δ 7.26 (s, 1H, =CH–), δ 3.40 – 3.42 (d, 2H, –CH_2_–, J= 6.9 Hz), δ 5.95 – 5.98 (m, 1H, =CH–), δ 5.10 – 5.12 (m, 2H, =CH_2_); ^13^C NMR (125 MHz, DMSO) d 169.1, 165.4, 156.7, 151.2, 137.7, 136.0, 135.4, 132.5, 122.4, 115.9, 113.5, 102.1, 100.9, 40.0; MS m/z: 284.3 [M]^+^

#### c. General procedure for preparation of compounds (7-8)

The 20 ml of NCS in Acetone: Water = 4:1 (1 mol) solution was put into the flask that was cooled in an ice-salt bath at −10^°^C before adding compound **(5)** and **(1)** (0.67 mol), respectively, and stirring it for 20 minutes. After 20 min of stirring, 30 L of cold water was added and the entire contents of the flask was transferred to a separatory funnel. The aqueous layer was removed, and the organic layer was washed with 20 ml of water, dried over with MgSO_4_, and filtered into a distilling flask [12]. The spectral characters of synthesized compounds **(7-8)** are as follows.

### 4-(2-chloro-3-hydroxypropyl)-2-hydroxyphenyl 2-hydroxybenzoate (7)

Yellowish liquid, Yield: 73.3%, FTIR: 3421 (–OH), 2884 (C-H, aliphatic), 1683 (C=O), 1415 (C=C aryl); ^1^H-NMR (500 MHz, DMSO-d6): δ 7.52 – 7.54 (d, 2H, =CH–, J= 8.0 Hz), δ 7.70 – 7.72 (m, 1H, =CH–), δ 8.09 – 8.10 (dd, 1H, =CH–, J= 8.0; 4.0 Hz), δ 7.03 – 7.05 (d, 2H, =CH–, J= 8.0 Hz), δ 7.09 – 7.11 (d, 1H, =CH–, J= 8.0 Hz), δ 2.80 – 2.82 (d, 2H, –CH2–, J= 7.0 Hz), δ 4.38 – 4.40 (m, 1H, –CH(Cl)–), δ 3.76 – 3.78 (m, 2H, –CH2–); ^13^C NMR (125 MHz, DMSO) d 168.5, 161.7, 151.5, 139.5, 136.7, 136.0, 130.5, 122.4, 119.3, 113.6, 111.8, 76.6, 70.0, 40.0; MS m/z: 322.0 [M]^+^

### 4-(2-chloro-3-hydroxypropyl)-2-methoxyphenyl 2-hydroxybenzoate (8)

Blackish brown powder, Yield: 57.6%, FTIR: 3409 (–OH), 2965 (C-H, aliphatic), 1660 (C=O), 1440 (C=C aryl), 1103 (C–O); ^1^H-NMR (500 MHz, DMSO-d6): δ 7.52 – 7.54 (d, 2H, =CH–, J= 8.0 Hz), δ 7.70 – 7.72 (m, 1H, =CH–), δ 8.09 – 8.10 (dd, 1H, =CH–, J= 8.0; 4.0 Hz), δ 7.03 – 7.05 (d, 2H, =CH–, J= 8.0 Hz), δ 7.09 – 7.11 (d, 1H, =CH–, J= 8.0 Hz), δ 2.80 – 2.82 (d, 2H, –CH2–, J= 7.0 Hz), δ 4.38 – 4.40 (m, 1H, –CH(Cl)–), δ 3.76 – 3.78 (m, 2H, –CH2–), δ 3.80 (s, 3H, –OCH_3_); ^13^C NMR (125 MHz, DMSO) d 168.5, 161.7, 151.5, 139.5, 136.7, 136.0, 130.5, 122.4, 119.3, 113.6, 111.8, 76.6, 70.0, 65.0, 40.0; MS m/z: 336.4 [M]^+^

#### d. General procedure for preparation of compounds (9-10)

Compound **(1)** (0,1 mol) was dissolved in mixing solvent (Ethanol:THF:H_2_O = 1:1:0,2). Then added (DHQ)_2_PHAL and (DHQD)2PHAL (0,01 mol), respectively, osmium catalyst (0,01 mol), and *N*-methylmorpholine oxide (NMO) (0,3 mol). The mixture was stirred for 2 hours at room temperature. The reaction was terminated by adding Na_2_SO_3_ and the mixture was extracted with CH_2_Cl_2_ (3×15 mL). The organic layer was dried with anhydrous MgSO_4_, filtered, and concentrated. The product was purified using chromatography column with appropriate mobile phase. The spectral characters of synthesized compounds **(9-10)** are as follows.

### 4-[(2S)-2,3-dihydroxypropyl]-2-methoxyphenyl 2-hydroxybenzoate (9)

Blackish white powder, Yield: 71.4%, FTIR: 3409 (–OH), 2965 (C-H, aliphatic), 1660 (C=O), 1440 (C=C aryl), 1103 (C–O); ^1^H-NMR (500 MHz, DMSO-d6): δ 7.52 – 7.54 (d, 2H, =CH–, J= 8.0 Hz), δ 7.70 – 7.72 (m, 1H, =CH–), δ 8.09 – 8.10 (dd, 1H, =CH–, J= 8.0; 4.0 Hz), δ 10.48 (s, 1H, –OH), δ 7.03 – 7.05 (d, 2H, =CH–, J= 8.0 Hz), δ 7.09 – 7.11 (d, 1H, =CH–, J= 8.0 Hz), δ 2.79 – 2.81 (d, 2H, –CH2–, J= 7.0 Hz), δ 4.21 – 4.23 (m, 1H, –CH(OH)–), δ 3.57 – 3.59 (m, 2H, –CH2–), δ 3.82 (s, 3H, –OCH_3_); ^13^C NMR (125 MHz, DMSO) d 168.5, 161.7, 151.5, 139.5, 136.7, 136.0, 130.5, 122.4, 119.3, 113.6, 111.8, 76.6, 68.8, 65.0, 40.0; MS m/z: 318.1 [M]^+^

### 4-[(2R)-2,3-dihydroxypropyl]-2-methoxyphenyl 2-hydroxybenzoate (10)

Blackish white powder, Yield: 51.8%, FTIR: 3326 (–OH), 2958 (C-H, aliphatic), 1685 (C=O), 1450 (C=C aryl), 1317 (C–O); ^1^H-NMR (500 MHz, DMSO-d6): δ 7.08 – 7.10 (d, 2H, =CH–, J= 8.0 Hz), δ 7.70 – 7.72 (m, 1H, =CH–), δ 8.09 – 8.10 (dd, 1H, =CH–, J= 8.0; 4.0 Hz), δ 10.48 (s, 1H, –OH), δ 6.86 – 6.87 (d, 2H, =CH–, J= 8.0 Hz), δ 7.03 – 7.04 (d, 1H, =CH–, J= 8.0 Hz), δ 2.78 – 2.80 (d, 2H, –CH2–, J= 7.0 Hz), δ 4.21 – 4.23 (m, 1H, –CH(OH)–), δ 3.71 – 3.73 (m, 2H, –CH2–), δ 3.82 (s, 3H, –OCH_3_); ^13^C NMR (125 MHz, DMSO) d 168.5, 161.7, 151.5, 139.5, 136.7, 136.0, 130.5, 122.4, 119.3, 113.6, 111.8, 76.6, 68.8, 65.0, 40.0; MS m/z: 318.1 [M]^+^

#### Purification and analysis of physicochemical properties

The result of the synthesis is seen by using the KLT method. Samples of synthesis are chromatographed with TLC to see the purity of the product. If the result is not pure, they are purified with recrystallization, separated by column chromatography method or other possible methods. Properties of pure compounds obtained are then analysed such as organoleptic, melting point, solubility in the solvent, refractive index and its calculated yield.

#### Cell culture

Cell cultures were taken from the stock stored in a liquid tank placed in the locator at a temperature of −196 °C. Cell cultures were thawed in water ± 37.7 °C. Then the cells were grown in some (2-3) small tissue culture flasks and incubated in incubator at speed of 37 °C with a flow of 5% CO_2_ and 95% O_2_. After 24 hours, the medium was replaced and the cells were grown until confluent and the amount was sufficient for research. After the sufficient number of cells or confluent (± 70%), the medium was replaced with a new medium RPMI 1640 as much as 5 ml. Cells were taken as much as 3 ⨯ 104 cells/100 μl of medium through the calculation done through the hemocytometer chamber [13].

#### Cytotoxicity test of MTT Method

The HT29 cell suspension of 100 μl with a density of 3 ⨯ 104 cells/100 μl of media was distributed into the 96-well plates and incubated for 24 hours. After incubation, the 100-μl test solution in various concentration series was poured into the plates. As a positive control, 100 μl Gossypol and Navitoclax in various series of concentrations were added into the plates containing 100 μl suspense cells. As the cell control, 100 μl culture media was added into the plates containing 100 μl of cell suspense and 100 μl DMSO was added into the plates containing 100 μl of cell suspension with delusions corresponding to the delusions of the test solution concentration as solvent control, then incubated for 24 hours in the incubator with a flow of 5% CO_2_ and 95% O_2_. At the end of incubation, culture media was discarded before added with 10 μl of MTT solution (5 mg/mL PBS), then the cell incubated for 3-4 hours. The MTT reaction was discontinued with the addition of SDS stopper reagent (100 μl). Microplate containing cell suspension was in shaken cell culture for ± 5 minutes then wrapped with aluminium foil and incubated for 1 night at room temperature. The living cells reacted to MTT forming a purple colour. Test result read with ELISA reader at 595 nm wavelength [14].

#### Molecular docking

Molecular docking experiment is performed using Autodock4 program (The Scripps Institute) to dock the eugenol derivatives to the binding site of the BCL-2 (PDB ID : 4LXD) using default parameters. Both protein and ligands are saved as output pdbqt files. For specific docking of ligand eugenyl benzoate derivatives onto the BCL-2 protein, the grid box volume was adjusted to 40×40×40 Å in the X, Y, and Z axes, respectively, with grid-sizes have a space up to 0.375 Å. Autodock4 will employs an idealized active site ligand as a target to generate putative poses of molecules. The binding energy values were calculated based on the total intermolecular energies (kJ/mol) including hydrogen bond energy, Van Der Waals energy, desolvation energy, and electrostatic energy. On the other hand, analysis of screening compounds was based on the energy variation, due to the formation of the ligand-receptor structure, it is given by the binding constant and the Gibbs free energy (ΔG) values. Prediction of the binding energy is performed by evaluating the most important physical-chemical phenomena involved in ligand-receptor binding, including conformation of the structure and hydrogen bonding interaction between compounds and the target protein.

## Results

### Synthesis of eugenol derivatives

In this work, we have successfully synthesized ten eugenyl-benzoate derivatives presented in the **Figure 1**. Once the synthesis is performed, the purification of the synthesized product was using column chromatography, with silica gel as stationary phase and appropriate mobile phase composition for each compound as seen in **Table 1**.

**Table 1.** Mobile phase for synthesized compounds purification using column chromatography

| Comp. Numb. | Compound | Mobile Phase |
| --- | --- | --- |
| (1) | 2-methoxy-4-(prop-2-en-1-yl)phenyl 2-hydroxybenzoate | Hexane : ethyl acetate 20 : 1 → 10 : 1 |
| (2) | 2-methoxy-4-(prop-2-en-1-yl)phenyl 3-amino-2-hydroxybenzoate | Hexane : ethyl acetate 20 : 1 → 10 : 1 |
| (3) | 2-methoxy-4-(prop-2-en-1-yl)phenyl 3,4,5-trihydroxybenzoate | Chloroform : methanol 50 : 1 |
| (4) | 2-methoxy-4-(prop-2-en-1-yl)phenyl 3,5-dihydroxy-4-methoxybenzoate | Chloroform : methanol 50 : 1 |
| (5) | 2-hydroxy-4-(prop-2-en-1-yl)phenyl 2-hydroxybenzoate | Hexane : ethyl acetate 20 : 1 → 10 : 1 |
| (6) | 2-hydroxy-4-(prop-2-en-1-yl)phenyl 3-amino-2-hydroxybenzoate | Hexane : ethyl acetate 20 : 1 → 10 : 1 |
| (7) | 4-(2-chloro-3-hydroxypropyl)-2-hydroxyphenyl 2-hydroxybenzoate | Hexane : ethyl acetate 20 : 1 → 5 : 1 |
| (8) | 4-(2-chloro-3-hydroxypropyl)-2-methoxyphenyl 2-hydroxybenzoate | Hexane : ethyl acetate 20 : 1 → 5 : 1 |
| (9) | 4-[(2S)-2,3-dihydroxypropyl]-2-methoxyphenyl 2-hydroxybenzoate | Hexane : ethyl acetate 10 : 1 → 1 : 1 |
| (10) | 4-[(2R)-2,3-dihydroxypropyl]-2-methoxyphenyl 2-hydroxybenzoate | Hexane : ethyl acetate 10 : 1 → 1 : 1 |

Preliminary analysis was done before the structure elucidation by using TLC at 254 nm wavelengths using UV light stain detector. Preliminary analysis results using TLC as listed are shown in **Table 2**. Based on the KLT test, it can be seen that the compound **(1)** is the most non-polar compound as it has the greatest Rf value among other compounds in the most non-polar eluent. Whereas the most polar compound is compound **(3)** as it has the smallest Rf value among other compounds in the most polar eluent. The presence of the demethylation reaction in the methoxy to hydroxyl group will also increase the polarity of the compound. As seen in compounds **(1)** and **(5)**, compound **(1)** has Rf value of 0.72 and compound **(5)** has Rf value of 0.36. It is influenced by the polar clusters –OH that exists in the compound. The more the number of hydroxyl groups, polarity of the compounds will be higher.

**Table 2.**
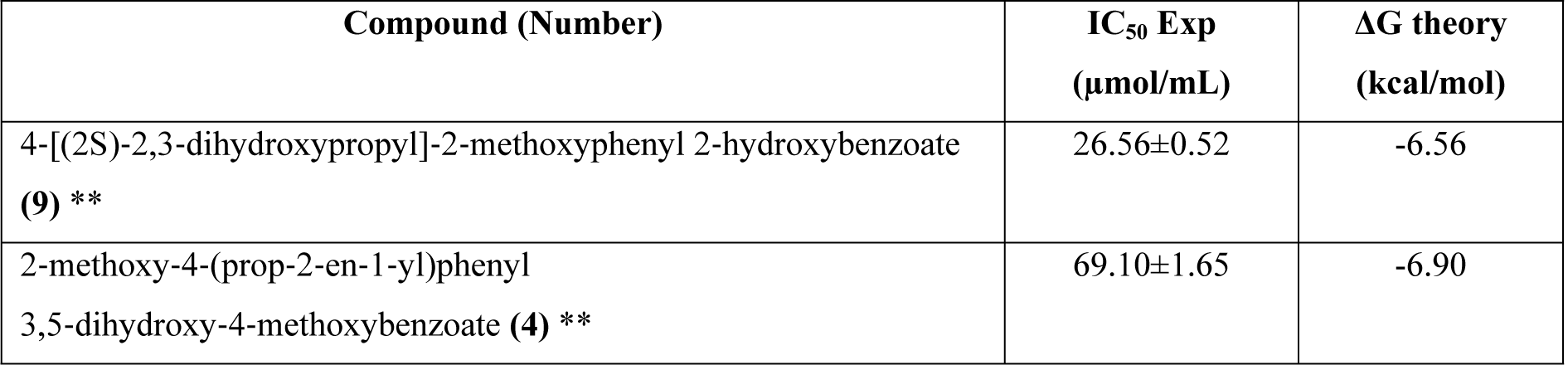
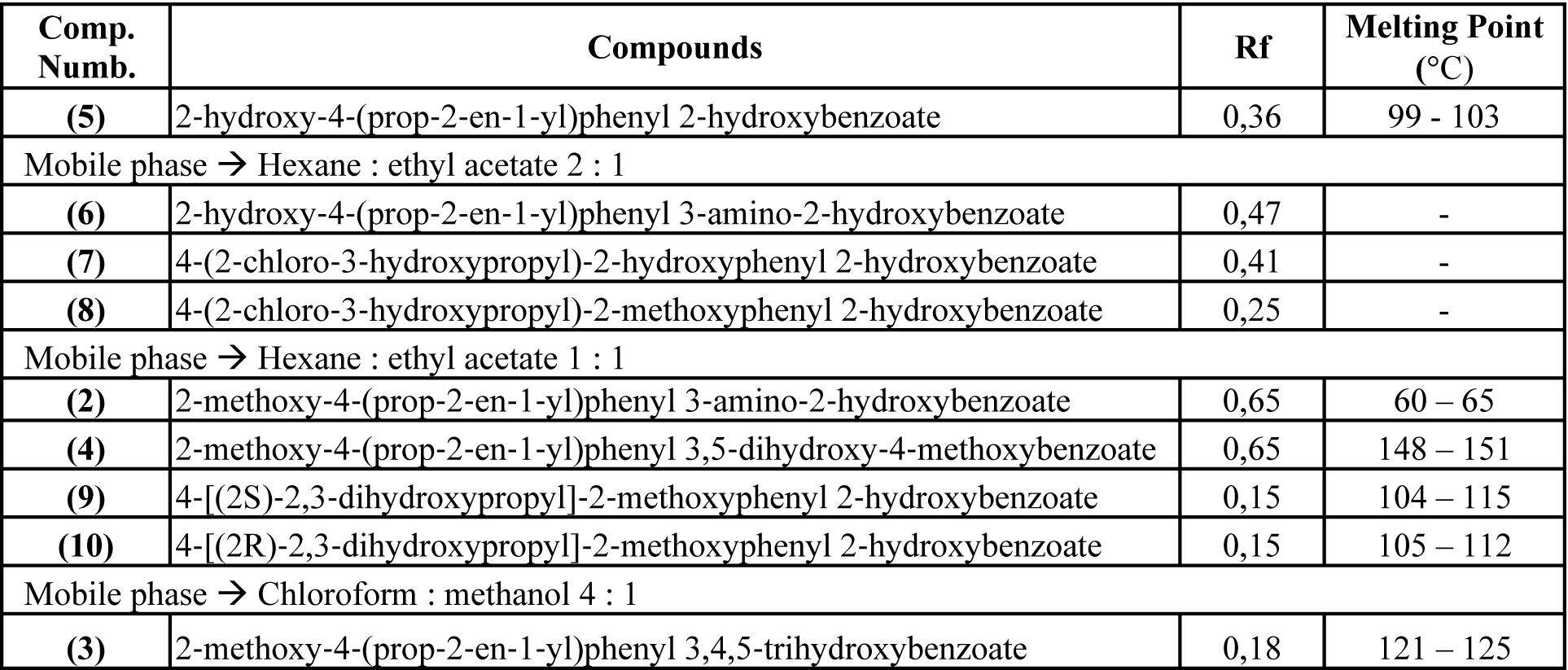
Physical data of the synthesized compounds

### Cytotoxicity Test of Eugenol Derivative Compounds

The results of the cytotoxicity of the compound as seen in **Table 3**, subsequently were processed by multiple linear regression analysis methods, by creating a graph stating the relationship between% inhibition with sample concentration log (μM). The graph shows data of linear equations and correlation (r). The IC_50_ value can be determined by inversing the y value = 50 on the chart to get the X value. The value of IC_50_ is an antilog from value X.

**Table 3.**
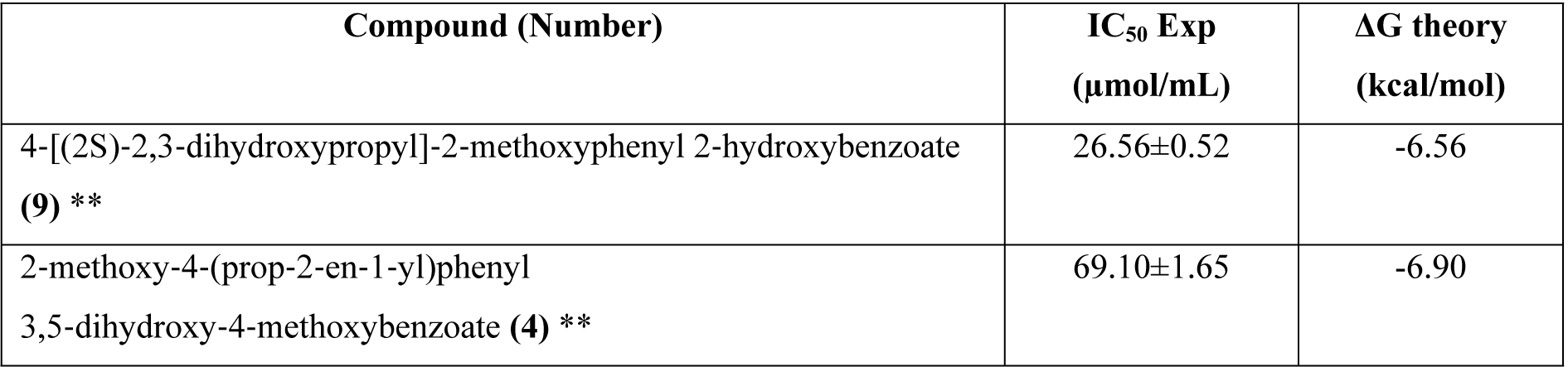
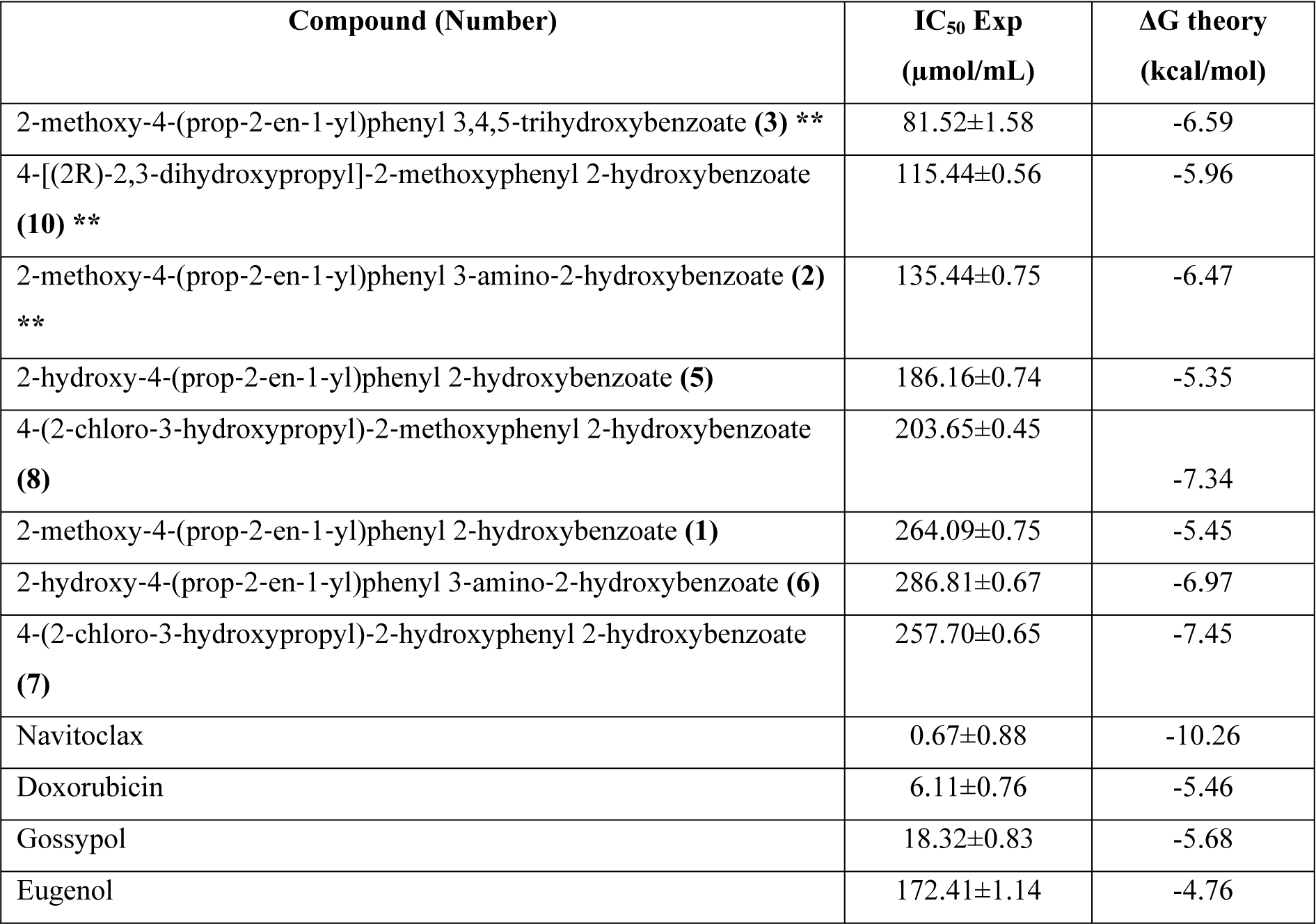
IC_50_ value of eugenol derivative compounds

In the **Table 3**, the value of IC_50_ for derivative compounds was between 26.56 μmol/ml −286.81 μmol/ml which the IC_50_ value of eugenol as lead compound was 172.41 μmol/ml. The greater the value of IC_50_ then the more non-toxic the compound is. The end of the cytotoxicity test on the target organ provides direct information about the changes occurring to the specific function of the cell.

### Quantitative structure-activity relationship and Molecular Docking

Analysis of Quantitative structure-activity relationship (QSAR) between eugenol derivative (**Figure 2**) with colorectal anticancer activity, particularly those utilizing computers based on theory by using calculation results. The electronic parameters used as a free variable in the QSAR analysis are the Molar Refraction (CMR) (steric parameters) and the logP for hydrophobic parameters (**Table 4**). MTT results indicate the IC_50_ value of the derivative compound is used as a bound variable. Analysis of QSAR was using multiple linear regression in SPSS.

**Table 4.**
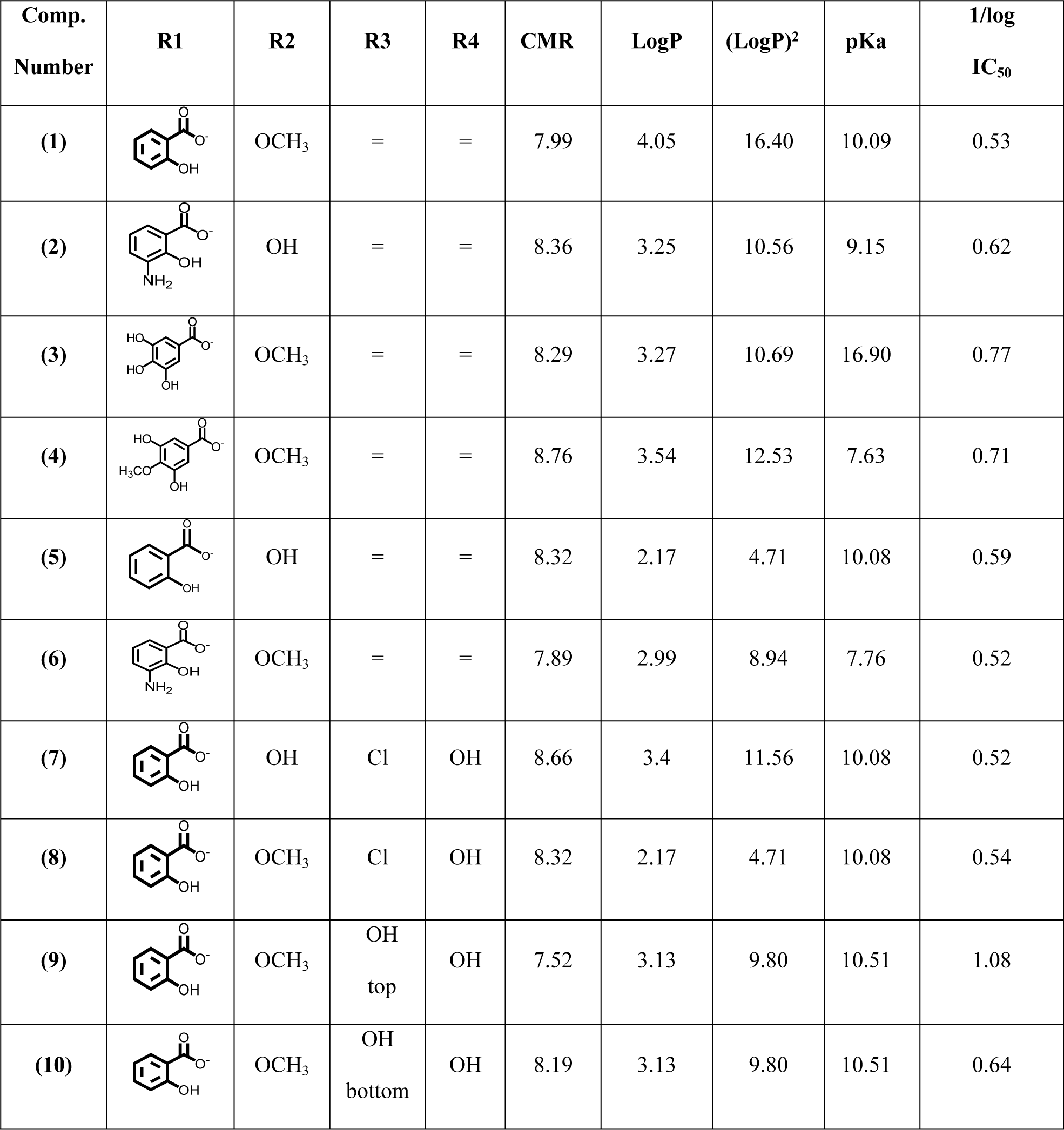
Analysis parameters of derivative compound structure activity

| Comp. Number | R1 | R2 | R3 | R4 | CMR | LogP | (LogP)<sup>2</sup> | pKa | 1/log IC<sub>50</sub> |
| --- | --- | --- | --- | --- | --- | --- | --- | --- | --- |
| (1) |  | OCH<sub>3</sub> | = | = | 7.99 | 4.05 | 16.40 | 10.09 | 0.53 |
| (2) |  | OH | = | = | 8.36 | 3.25 | 10.56 | 9.15 | 0.62 |
| (3) |  | OCH<sub>3</sub> | = | = | 8.29 | 3.27 | 10.69 | 16.90 | 0.77 |
| (4) |  | OCH<sub>3</sub> | = | = | 8.76 | 3.54 | 12.53 | 7.63 | 0.71 |
| (5) |  | OH | = | = | 8.32 | 2.17 | 4.71 | 10.08 | 0.59 |
| (6) |  | OCH<sub>3</sub> | = | = | 7.89 | 2.99 | 8.94 | 7.76 | 0.52 |
| (7) |  | OH | Cl | OH | 8.66 | 3.4 | 11.56 | 10.08 | 0.52 |
| (8) |  | OCH<sub>3</sub> | Cl | OH | 8.32 | 2.17 | 4.71 | 10.08 | 0.54 |
| (9) |  | OCH<sub>3</sub> | OH<br>top | OH | 7.52 | 3.13 | 9.80 | 10.51 | 1.08 |
| (10) |  | OCH<sub>3</sub> | OH<br>bottom | OH | 8.19 | 3.13 | 9.80 | 10.51 | 0.64 |

**Figure 2.**
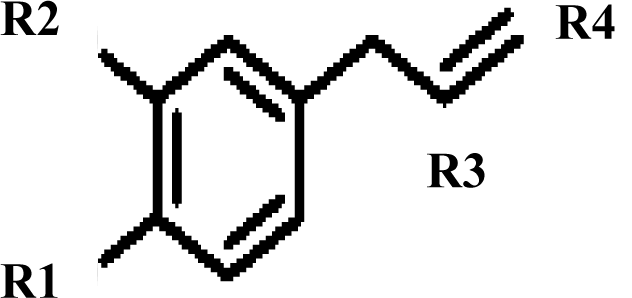
Structure modification of QSAR

The results of QSAR by multiple linear regression equations involving the properties of hydrophobicity (logP) and steric (CMR) parameters as inhibition of HT29 colorectal cancer cells as follows:

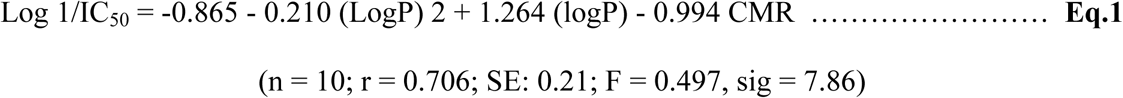

From the experimental data of eugenol derivatives, it can be concluded that there is a meaningful non-linear (parabolic) relationship between the hydrophobicity parameter (logP) of the eugenol derivatives with activity as HT29 colorectal cancer cell inhibitor; there is non-linear (parabolic) and meaningful relationship between the steric parameter of the eugenol derivative compounds (CMR) toward the inhibition activity of HT29 colorectal cancer cells; and the hydrophobicity parameter (logP) plays more role than the steric (CMR) ones.

## Discussions

In this work, we have successfully synthesized ten eugenyl-benzoate derivatives. All of the ten compounds structure were analysed using various spectroscopic analysis. The ^1^H-NMR for compounds **(5)** and **(6)** showed disappearances of singlet of –OCH_3_ around δ 3.80 ppm due to demethylation reaction of compounds **(1)** and **(2)**, respectively. Due to Halohydrin and Sharpless reaction, the ^1^H-NMR for compound **(7 – 10)** showed shifting from double bond of eugenyl structure (compound **(1-6)**) to single bond. Compounds with double bond **(1-6)** showed multiplet signal of =CH– around δ 6.00 ppm and multiplet signal of =CH_2_ around δ 5.15 ppm. Compounds with single bond **(7 – 10)** showed multiplet signal of –CH(Cl / OH)– around δ 4.2 – 4.3 ppm and multiplet signal of –CH_2_– around δ 3.5 – 3.7 ppm due to presence of –Cl or –OH.

From the QSAR analyses, it showed that the partition coefficient (logP) is logarithmically related to free energy. The partition coefficient logarithm as a parameter indicates that the steric and hydrophobic effects should be fully optimized for better BCL-2 inhibitory activity. Compounds that have low solubility in water are not able to penetrate the hydrophilic barrier and vice versa. The relationship between biological activity and the hydrophilic-lipophilic properties can be expressed with logP, where the solubility of the compound in certain conditions will be optimum and is described as a parabolic curve (logP)^2^.

The **Equation 1** shows that fewer substituents and less hydrophobicity in the main structure are very beneficial for selective enzymes and good biological activity as seen from the methoxy derivative group (less polar) on R2. Compounds with electron-rich halogen groups are lipophilic and able to penetrate the lipid bilayer of the target compound, as well as participate in molecular interactions that contribute to protein ligands binding [6]. In R3 position, the electron extractor like Cl severely undermines the inhibitory activity of BCL-2. Meanwhile, an addition modification in the double-bond group of eugenol is performed by hydrolysis and halohydrin on R3 and R4. Modifications involving the symmetry CH as in the compounds **(9)** and **(10)** indicate the presence of the bottom up position and top up of the hydroxyl position in R3 definitely affects the activity of derivative compounds. On R3 – OH bottom up position has the lowest IC_50_ value, it proves that a symmetry CH factor affects the inhibitory activity against HT29 colorectal cancer cells.

The analysis of the docking results in **Figure 3** indicates that the **(9)** compound has hydrogen binding with BCL-2 in the Arg143 and Ala146 positions where both of these amino acids are in binding site ligands of the BCL-2. Because of that, **(9)** is more stable than other derivatives. In line with research conducted by Fukushi *et al*. (2014), modification using addition reaction of the double-bond of benzoate in the terminal hydroxyl group proved to have anticancer activity. This increased activity is due to a terminal hydroxyl group that can increase the coefficient of octanol-water partitions and the topological polar surface area (TPSA). This study provides a better insight into the formulation of a BCL-2 inhibitor in the future before its synthesis.

**Figure 3.**
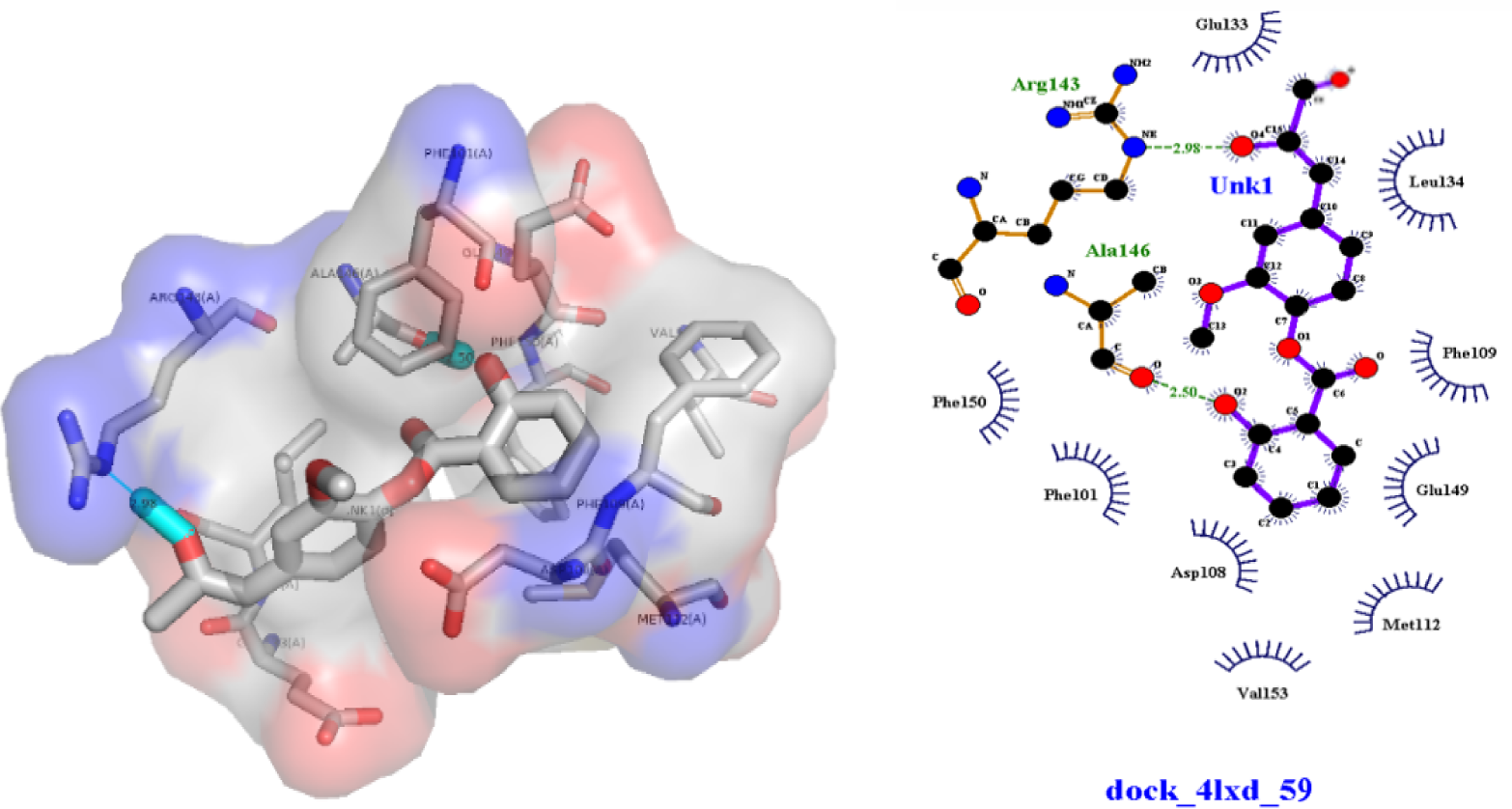
Interaction Hydrogen bond of eugenyl benzoate derivatives with BCL-2

## Conclusions

The synthesis of new compounds was done with Mitsunobu’s esterification reaction and then continued with Sharpless reaction, produced the best active compound **(9)** as BCL-2 inhibitors better than other eugenol derivatives. QSAR indicates the logP and CMR have effect on its colorectal cytotoxic activity which the hydrophobicity parameter (logP) plays more role than the steric parameter (CMR).

## Acknowledgements

The authors acknowledge the financial support received from Universitas Indonesia PUTI Q1 Grant year 2020 (Grant number: NKB-1297/UN2.RST/HKP.05.00/2020).

## Authors’ Contributions

Concept and design: FF, RA, AA, AY; Data acquisition: FF, RIP; Data analysis: FF, RIP, Drafting manuscript: FF, RIP, LE; Critical revision of manuscript: FF, RIP; Statistical analysis: FF, LE; Supervision: RA, AA, AY. All authors read and approved the final manuscript.

